# A spiking neural circuit model for learning multi-sensory integration

**DOI:** 10.1101/2020.11.27.401216

**Authors:** Deying Song, Xueyan Niu, Wen-Hao Zhang, Tai Sing Lee

**Affiliations:** Yuanpei College, Peking University; Neural Science PhD Program, New York University; Department of Mathematics, University of Pittsburgh; Center for the Neural Basis of Cognition, Carnegie Mellon University

## Abstract

Neurons in visual and vestibular information integration areas of macaque brain such as medial superior temporal (MSTd) and ventral intraparietal (VIP) have been classified into congruent neurons and opposite neurons, which prefer congruent inputs and opposite inputs from the two sensory modalities, respectively. In this work, we propose a mechanistic spiking neural model that can account for the emergence of congruent and opposite neurons and their interactions in a neural circuit for multi-sensory integration. The spiking neural circuit model is adopted from an established model for the circuits of the primary visual cortex with little changes in parameters. The network can learn, based on the basic Hebbian learning principle, the correct topological organization and behaviors of the congruent and opposite neurons that have been proposed to play a role in multi-sensory integration. This work explore the constraints and the conditions that lead to the development of a proposed neural circuit for cue integration. It also demonstrates that such neural circuit might indeed be a canonical circuit shared by computations in many cortical areas.

## 1 Introduction

Traditionally, researchers have focused on studying an individual sensory modality, like vision, audition and touch, in isolation. Although this has undoubtedly provided us with great amounts of knowledge about the basic mechanisms underlying information processing and unimodal attention, it is unsatisfactory when it comes to our rich multisensory experiences and behaviors. [1] There have been numerous neurophysiological and behavioral studies demonstrating extensive interactions among senses, as well as researches indicating the processes underlying multisensory information processing. [2]

In this study, we focus on information integration between visual inputs and vestibular inputs. They both convey information about heading direction. When we walk on the street, the optical flows we see and the vestibular signals we experience are in consistence. This is when an integration model will be selected to present more faithful estimate of heading direction of self-motion. However, when we wear a virtual reality headset and sit still on a chair, visual and vestibular signals are discordant and this is the case where a segregation model should be used.

Whats the neural implementation of visual and vestibular integration? There are two kinds of multisensory neurons in visual and vestibular brain areas, such as ventral intraparietal (VIP) areas and dorsal medial superior temporal area (MSTd). They are named according to whether their tunings under each sensory cue are congruent or opposite. Congruent neurons prefer visual and vestibular cues of the same direction while opposite neurons prefer cues of opposite directions. [3] [4] [5] [6] [7]

Previously, a firing-rate model has been proposed to achieve visual and vestibular sensory integration. [8] [9] But the congruent neurons in that model could send both excitatory and inhibitory signals to other neurons, violating Dales law. Also their normalization were done by activation function. In this work, we adopt a more detailed biophysical model which transfers the firing rate model to a spiking model with both excitatory and inhibitory neurons to achieve multisensory integration. Inhibitory pools of neurons are introduced not only for normalization, but also for interaction with opposite neurons. The learned neurons have tuning properties agreeing with experiments as well as theoretical predictions. We have also discussed about parameter regime and their implications, which provides more detailed information about the stability and sensitivity of the model.

In the following sections, we are going to use a spiking network inspired by a well-established V1 circuit [10] [11] to learn the feedforward connections to congruent and opposite neurons and also the horizontal connections between two multisensory integration modules. We are also going to show the learned netwrok behaves as expected in the sense that the population response and single neuron tuning curve are consistent with experiments as well as theoretical predictions.

## 2 Materials and Methods

### 2.1 Model setup

Figure 1 indicates the model architecture. Red circles indicate excitatory neurons while blue circles indicate inhibitory neurons. Arrow indicates excitatory connections while a dot indicates inhibitory connections. Each module is mainly composed of a pool of sensory neurons, a pool of congruent neurons and a pool of opposite neurons. I11, I13, I21, I23 are for normalization. Normalization not only makes our model more biologically plausible, but also is necessary for our model to work properly, as seen in later sections. I12 and I22 are inhibitory interneurons implementing the inhibition from C to S without violating Dales law.

**Figure 1:**
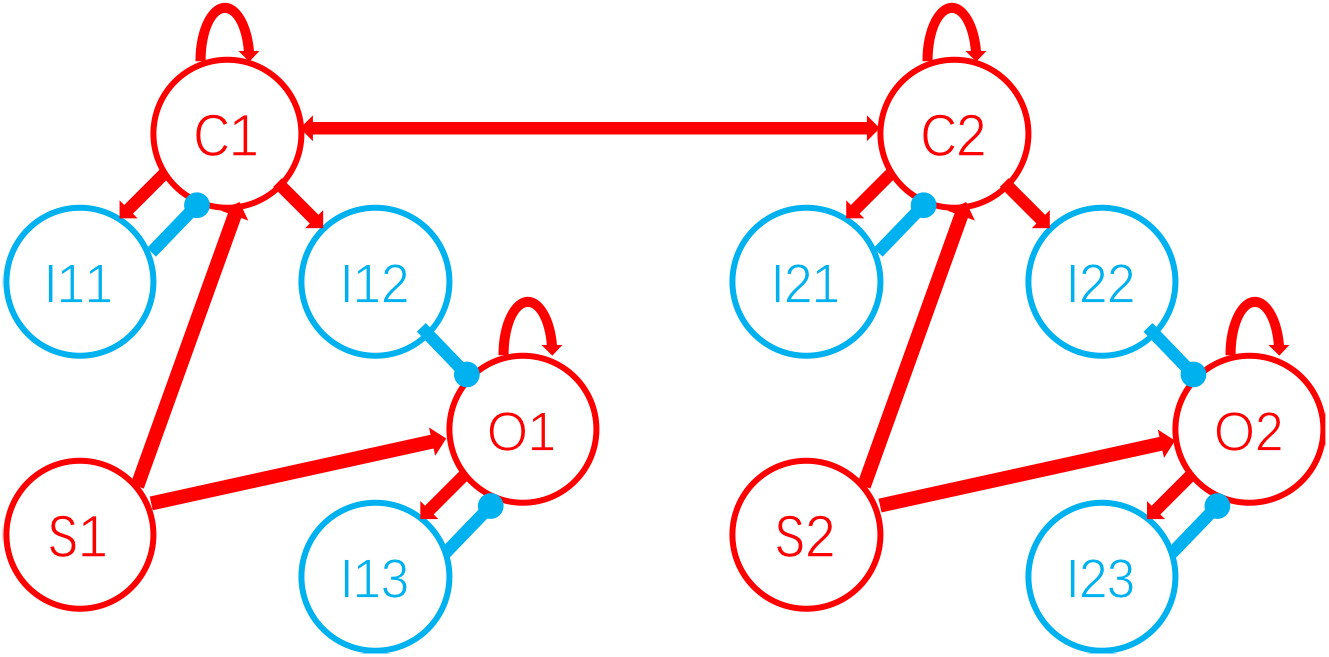
Model architecture.

There are recurrent connections among each pool of congruent and opposite neurons. There are also excitatory reciprocal connections between C1 and C2 bridging the two modules.

There are 36 neurons in each of the excitatory ring and I12, I22, 9 neurons in I11, I13, I21, I23. Each neurons dynamic is governed by a leaky integrate and fire function.

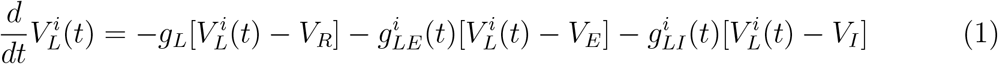

where *L* indicates which pool (ring) the neuron belongs to, and *i* is the index of the neuron in that pool. The functions 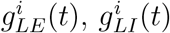 denote the temporal profiles of the excitatory (inhibitory) conductance that impinge upon the *i^th^* neuron in pool *L*. When the voltage 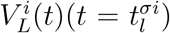 reaches spiking threshold *V*_*T*_, the spike time 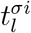 is recorded, the voltage is reset to *V*_*R*_, held there for a brief time *t*_*R*_, and then re-initialized as 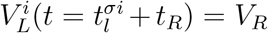 “leak conductance” *g*_*L*_, the reversal potentials (*V*_*R*_, *V*_*E*_, *V*_*I*_) are constants, as is the threshold *V*_*T*_. The values for the biophysical parameters are commonly accepted. The “refractory period” *t*_*R*_ = 3*ms*(1*ms*) for excitatory (inhibitory) neurons. After nondimensionalization, *g*_*L*_ = 50, *V*_*I*_ = − 2*/*3, *V*_*R*_ = 0, *V*_*T*_ = 1, *V*_*E*_ = 14*/*3.

The conductances in equation (1) drives the time evolution of the membrane voltage.

For I,

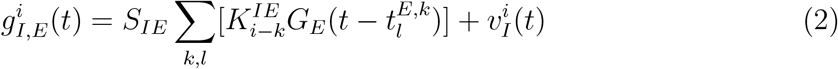

For C1,

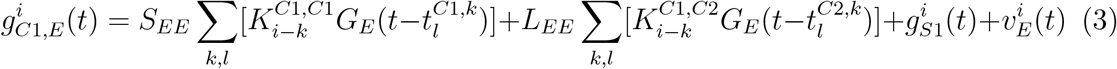

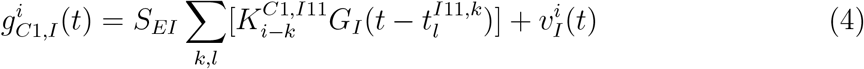

For O1,

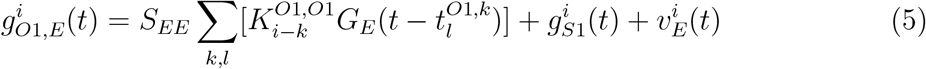

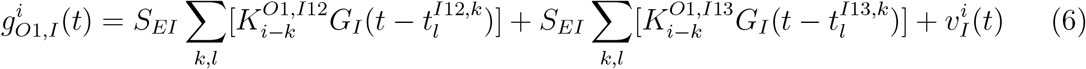

In (2)–(6), *S*_*σ*_1__, _*σ*_2__ is synaptic strength, where *σ*_2_ is the sending neuron type and *σ*_1_ is the receiving neuron type. *k* indexes the number of sending neurons in its pool, while *l* indexes the number of spike of that particular neuron. As for the spatial kernels, they are von Mises.

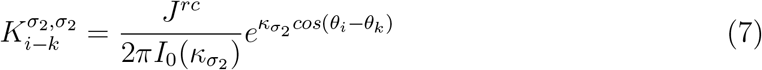

for pools of neurons in the same module, and

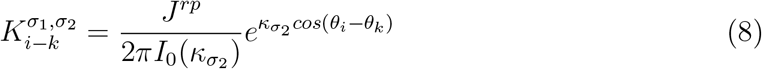

for pools of neurons in different modules. *I*_0_(*x*) is the modified Bessel function of the first kind with order 0. However, the interactions between the excitatory pool and normalization pool are uniform. That is,

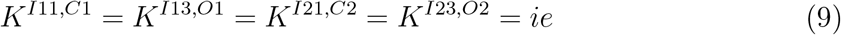

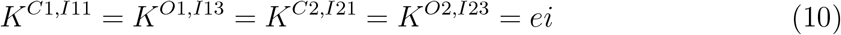

We need to mention here that the spatial kernels *K*^*C*1,*S*1^,*K*^*O*1,*S*1^, *K*^*O*1,*I*12^, *K*^*C*2,*C*1^,*K*^*C*2,*S*2^, *K*^*O*2,*S*2^, *K*^*C*2,*I*22^, *K*^*C*1,*C*2^ are learned.

As for temporal kernels, *G*_*E*_ is sum of two alpha functions for AMPA and NMDA respectively, while *G*_*I*_ is sum of two alpha functions for *GABA*_*A*_ and *GABA*_*B*_ respectively. Alpha function can be written as

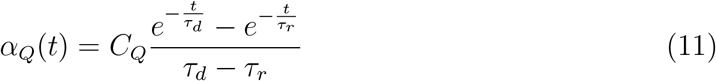

When *τ*_*d*_ = *τ*_*r*_,

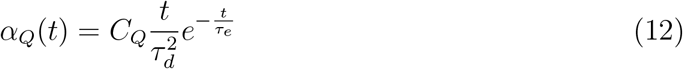

where Q indexes different neurotransmitters. The S conductance in the equation is the sum of synaptic conductance waveforms evoked by S spikes, with kinetics like the conductance waveforms of that in C, I and O. Each S neuron’s spike train is modeled as a modulated Poisson process. The modulation of the spike rate of the *i^th^* S neuron is given by

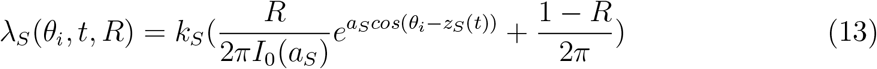

The subscript *S* ∈ {1, 2} indicates whether the input is from S1 or S2. *k*_*S*_ is a scaling constant, while *R* ∈ [0, 1] is the reliability of the input. *a*_*S*_ determines the width of the input, while *z*_*S*_(*t*) refers to the center of the input at time t. Finally, 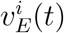 and 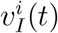are noise terms representing inputs from other cortical area,

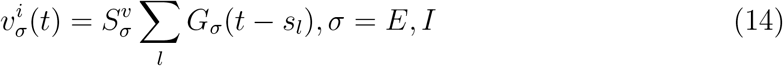

while the spike times *s*_*l*_ is a Poisson process with a fixed rate.

### 2.2 Learning algorithm

The feedforwarrd weights are all initialized with the following metohd: consider feedforward connections from a pool of input neurons indexed by *j* to a pool of target neurons indexed by *i*. For each *j*, we sample 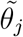 from a uniform distribution over the N target neurons without replacement (i.e., 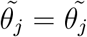, if and only if*j* = *j’*), as well as the multiplicative factor *A*_*j*_ from a log normal distribution with arithmetic mean of 0.3 and arithmetic variance of 0.1. Then the initial feedforward connections are given by

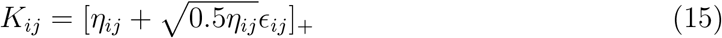

where

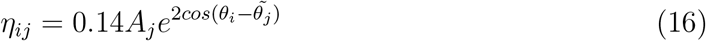

and *ϵ*_*ij*_ is i.i.d. Gaussian noise with *µ* = 0 and *σ* = 0.1. Intuitively, this models each input neuron as projecting to a random target location with variable connection strength and a spatial spread given by von Mises distribution.

The position of input from S1, *z*_1_(*t*), is generated by first randomly permuting an evenly spaced sequence of input from *π* to *π*, each lasting 2s. The simulation is run for 3600s and the step size for membrane potential update is 0.1ms. The weights are updated every 50ms, with time window being 100ms. Weights adjusted with the rule are further constrained to be non-negative.

We adopt Willshaw’s rule. It can be written as

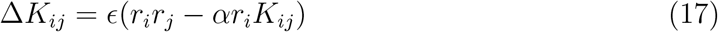

Learning rate *ϵ* are different for different set of connections.

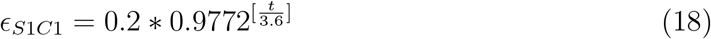

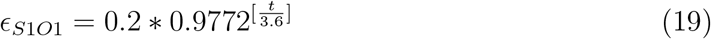

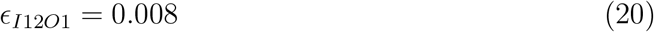

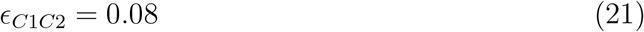

We adopt a curriculum learning method. Except Wi12o1, other connections starts to be learned at 0s. Wi12o1 starts to be learned at 2400s.

## 3 Results

### 3.1 Learned feedforward connections

The network exhibits self-organization, i.e., neurons in each ring learn to be topographically organized unsupervisedly. Topographically organized means that the neurons are arranged in a structure where there is an orderly spatial relationship between the distribution of sending neurons and a related distribution of receiving neurons.

According to Figure 1, we call the connections from Si to Ci, Si to Oi, and Ii2 to Oi feedforward connections, where i = 1 or 2 representing the two modules. The connections from Si to Ci, from Si to Oi are called direct feedforward connections, for they represent feedforward input directly from sensory neurons. We call the connections within congruent neurons and opposite neurons recurrent connections, and call the bi-directional connections between C1 and C2 reciprocal connections.

Only the connections mentioned above needs to be learned. It is natural that we dont need to learn normalization and recurrent connections. Also, we assume that WCiIi2 is arranged according to their spatial location, i.e., a von Mises connection from Ci to Ii2. The reason we introduce 6 pools of inhibitory neurons is to implement the function of congruent and opposite neurons without violating Dales law.

Figure 2A illustrates the connection from S1 to C1. Different colors indicate different preferred direction in S1 and the x-axis is by preferred direction in C1. It can be observed that the shape of feedforward connections is approximately a von-Mises distribution. It is usually assumed that feedforward connections are von-Mises or Gaussian shaped in multisensory integration models. We show here that a von Mises shape can naturally appear in our learning model. Figure B and C are similar. We only show results of the first module, for the two modules are symmetric in the model.

**Figure 2:**
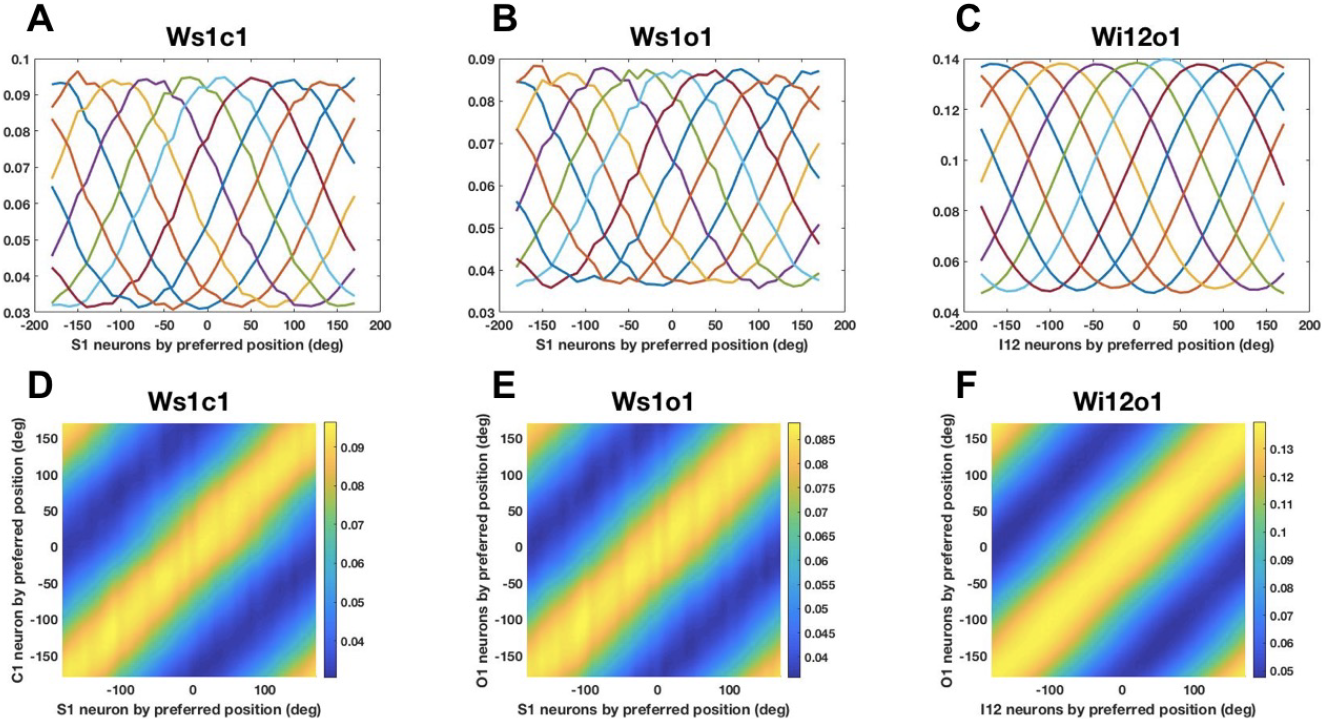
Learned feedforward connections and their topological order. A-C) feedforward connections in module 1. Different colors indicate neurons preferring different direction in the sendng pool. X-axis is by preferred direction in the receiving pool. D-F) The color indicates strength of connection. These figures illustrate that topological order is preserved for there is a bright band in the diagonal showing that all pairs of neurons preferring the same direction are strongly wired together. This is due to recurrent connections inside each individual ring.

**Figure 3:**
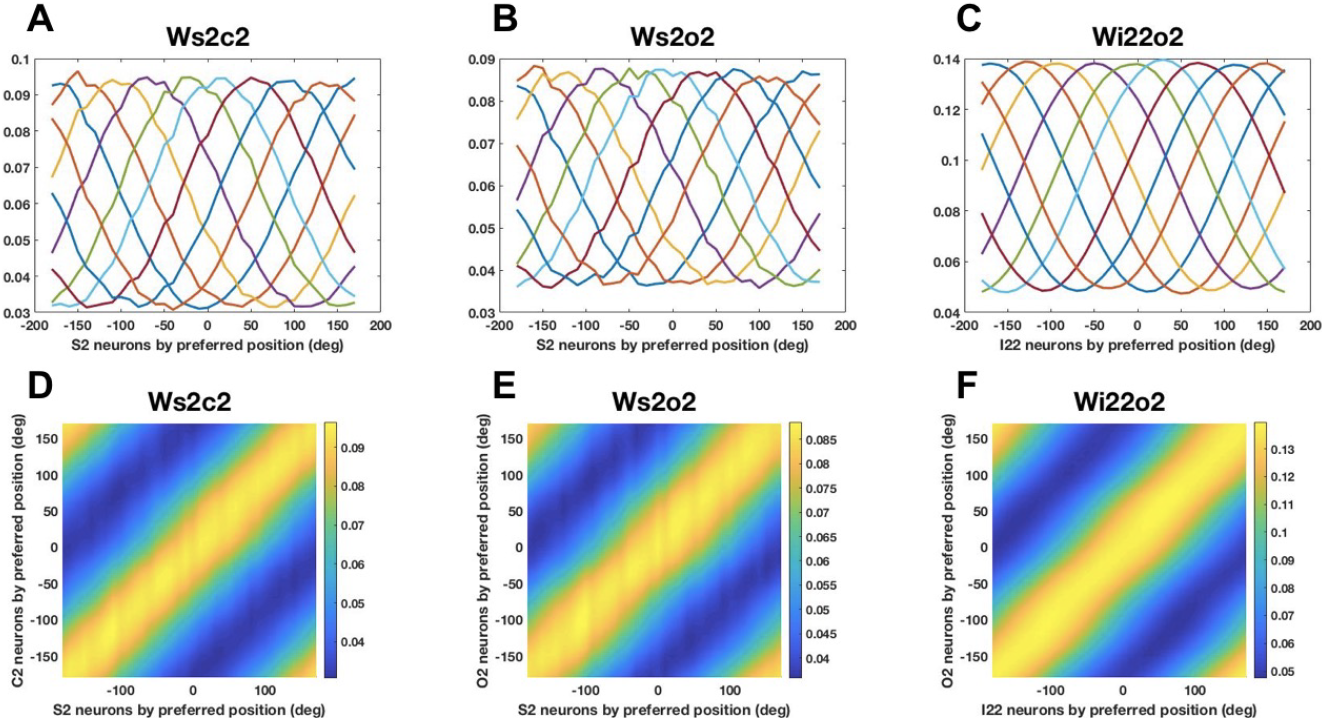
Learned feedforward connections and their topological order. Same as Figure 2, but for that in module 2.

In figure 2D-F, we show that the topological order is preserved from sensory neurons to congruent neurons, from congruent neurons to opposite neurons and from sensory neurons to opposite neurons. In all three figures, there is a bright diagonal indicating that all pairs of neurons preferring the same direction are strongly connected together. This phenomenon comes out from the fact that in each ring, the closer the neurons preferred directions are, the stronger the recurrent connection between them is. To some extent, the recurrent connection plays the role of teacher teaching the feedforward connections to arrange themselves.

### 3.2 Learned reciprocal connections

There is no surprise that the reciprocal connections between congruent neurons bridging the two modules also have von-Mises shape. Although the kernel is roughly of the same magnitude with that of the feedforward connections, the scale factor before the kernel of reciprocal connections is actually much smaller than that of feedforward connections. This is due to preventing the congruent neurons from spontaneous activities. Therefore, the information from the other module is much smaller than information from the self module, which will be demonstrated in the population response. As illustrated in Figure 4C-D, Wc1c2 and Wc2c1 are also topologically organized.

**Figure 4:**
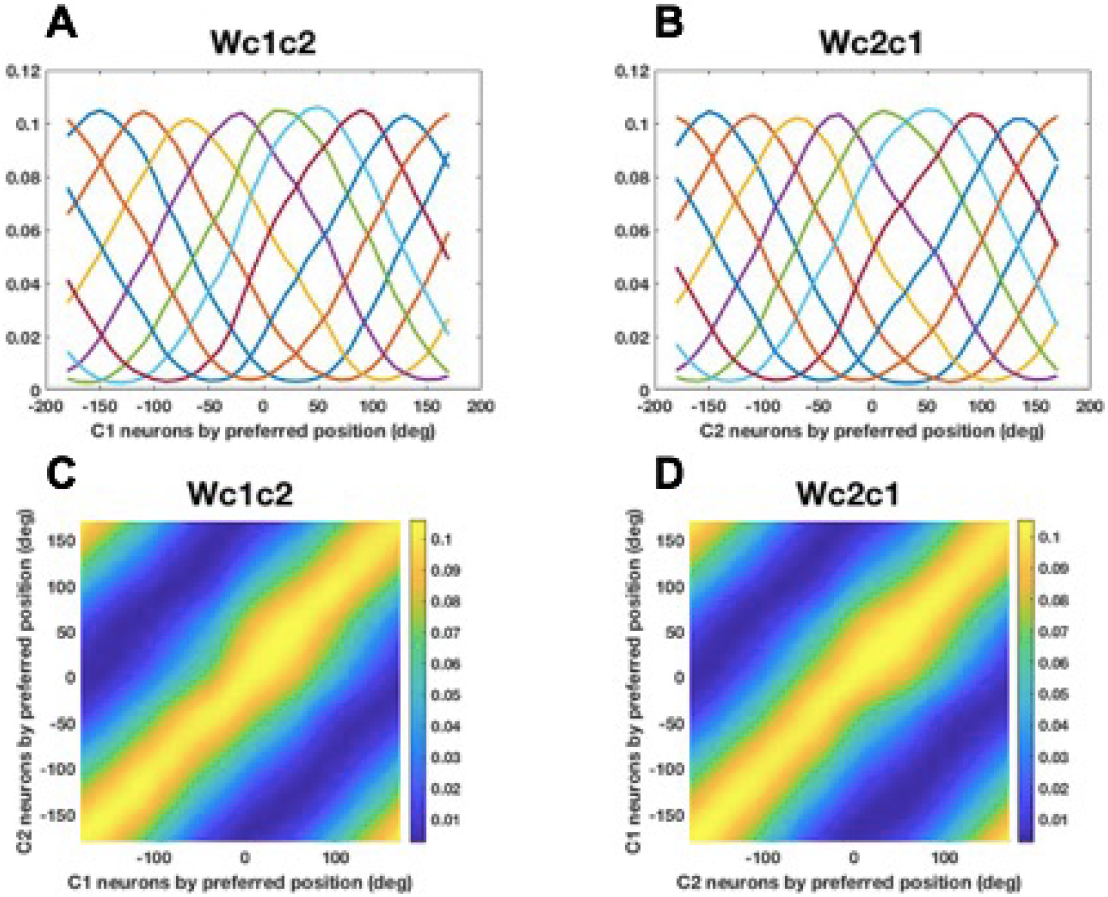
Learned reciprocal connections between two modules and their topological order. A) Reciprocal connections from module 1 to module 2. B) Reciprocal connections from module 2 to module 1. C-D) Topological order of neurons bridging the two modules.

### 3.3 Population response with learned connection weights

For the learned model, the population response of neurons in C1, O1, C2, O2 is in Figure 5 below. S1 is centered at 0 degree, and S2 is centered at −120 degree. We will see why congruent neurons are called congruent and opposite neurons opposite. Actually, the population response of congruent neurons always peaks in congruence with its unimodal stimulus, i.e., S1 or S2. As for opposite neurons, they tend to peak in consistence with input in their own module, while opposite to that from the other module.

**Figure 5:**
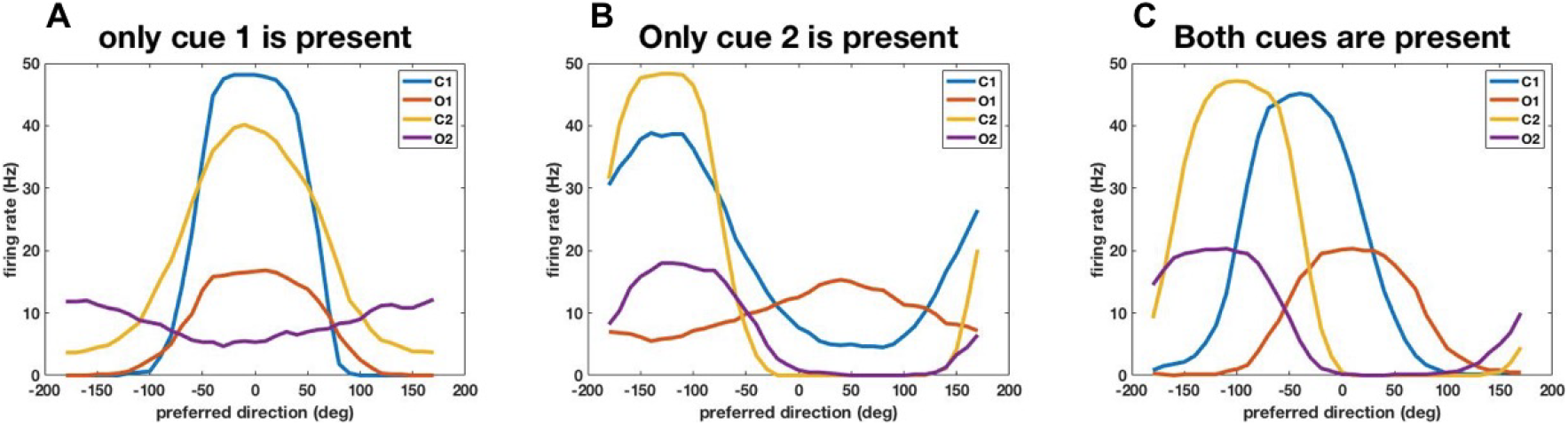
Population response of neurons in C1, O1, C2, O2. A) Only cue 1 is present. A) Only cue 2 is present. C) Both cues are present.

When only cue 1 is present, C1 will certainly peak at 0 degree. O1 will also peak at 0 degree for there is no input in module 2. It receives feedforward excitation from S1 and horizontal inhibition from C1. They both peak at 0 degree but the feedforward excitation is much stronger, so O1 will still peak at 0 degree. C2 will peak at 0 degree but with a smaller amplitude and a DC shift. The smaller amplitude is due to weaker input strength from C1 than S1 and The DC shift is due to input from S2 with zero reliability. O2 will peak at 180 degree, which is opposite to the center of S1. It receives DC input from S2 and horizontal inhibition from C2. C1 neuron will inhibit the most the neuron in S1 which has the same orientation preference. So neuron favoring 0 degree in O2 will receive the strongest inhibition, thus making the population response of O2 center at 180 degree, opposite to 0 degree.

When both cues are present, the peak of the response of C neurons lies between the center of two cues, but closer to the one in its own module. This can be understood intuitively for the input from the other module is indirect and thus weaker. The opposite neurons reflect the disparity of the two stimuli. For example, the peak the population response of O2 neurons lies between S2 and the opposite of S1 (S1+180 degree), but it’s closer to S2.

### 3.4 Single neuron response

We simulated the neurophysiological experiments presenting stimuli centered at different locations and measuring the response of a particular neuron to those stimuli. Here, ‘only cue 1 is present’ means that the reliability of cue 2 is zero instead of the input firing rate of S2 being zero, that is, in the formula of *λ*, *R*1 = 1 and *R*2 = 0. Note that there is still input in S2 even in the case of unimodal S1 stimulus, though the input in S2 is uniform across all directions. This is consistent with the observation that MT neurons appear to have s non-zero background input. What’s more, we mention here that a certain kind of homeostasis has to be maintained for our model to work: the total input from S1 and S2 has to remain relatively constant. This necessitates the use of a constant DC input in S2 even in the case of unimodal S1 stimulus.

Fig. 7A shows the tuning curve of a neuron in C1 preferring stimulus of −90 degree. When only cue 1 is present, the response of the neuron peaks at −90 degree with a von-Mise shape. When only cue 2 is present, the neuron’s response still peaks at −90 degree but the amplitude is less. This is due to the input from module 2 is indirect and relatively weaker. When both cues are present, the tuning of the congruent neuron has a similar von-Mise shape with stronger yet sub-additive response. Fig. 7B show the tuning curve of a neuron in O1 preferring −90 degree in module 1 and 90 degree in module 2. When only cue 1 is present, the response of the neuron peaks at −90 with a von-Mise shape but with smaller amplitude than that of C1. This is because besides excitation from S1, O1 also receives inhibition from C1. When only cue 2 is present, the response of the opposite neuron peaks at 90 degree. When both cues are present, the response is flattened and highly sub-additive. The subadditivity of the multisensory neurons’ response is consistent with experimental observations of MSTd neurons in macaque.

Fig. 7C-D show the correlation of congruent and opposite neurons’ tuning curves towards unimodal S1 stimulus and unimodal S2 stimulus, showing they have congruent and opposite tuning properties respectively. Responses to S1 and S2 stimuli (unimodal von-Mise) are most strongly correlated when the inputs are at the same location for congruent neurons. While responses are most strongly correlated with the inputs biased by 180 degree for opposite neurons.

## 4 Discussion

This work provides a biological implementation of a previous model with inhibitory neurons implementing normalization explicitly and the anti-Hebbian connection between C1 and O1 neurons. In this work, no a priori assumption about topographic organization of the multisensory neurons is made. The topographic organization naturally emerges via a Kohonen map-like mechanism.

We adopt a biologically realistic rate-based model to learn both congruent and opposite neurons. The learned opposite neurons exhibit experimentally observed tuning properties to bimodal stimuli and are topographically organized. Population response of congruent and opposite neurons are also qualitatively consistent with experimental results. Our model architecture is compatible with some existing decentralized models of multisensory integration, and therefore our work also provides a basis for leanring such models in general. Also, this network is inspired by and transferred from a model that was developed for V1, indicating that this model might be a biologically realistic canonical model.

### 4.1 Delicate balance between excitation and inhibitory normalization

We once adopted von-Mises excitatory to inhibitory and inhibitory to excitatory connection. However, there was always a two-period issue, i.e., there are two instead of one bright band after reordering in the topological order diagram. Thus, we turned to uniform inhibition between excitatory pool and inhibitory normalization pool instead of von-Mises. Then the issue was solved.

Intuitively, it is important for each stimulus, there is a clear winner, so that it can suppress the others. A von-Mises inhibition kernel tends to suppress oneself, and if the width is not wide enough, it will allow another winner to emerge at the far away, causing the pool to learn two cycles in the topology, two sets of neurons with the same receptive field. That is to say, there has to be an intricate balance between excitation and inhibitory normalization.

We then explored the parameter space more carefully, finding the parameter range that works is quite narrow. We first ask what the situation will be in some extreme values, which is when the inhibition is too weak or too strong. According to Figure 6A, when the inhibition is not strong enough, there is no clear winner to be leader for each stimulus, then everyone just follows the fashion no stability or everyone becomes the winner equal opportunity to learn. According to Figure 7B, when the inhibition is too strong, there will be no winner and the response is quite small, thus learning messy topological orders. It has to be mentioned that both results in Figure 6 are learned with constant learning rate. This is because we want to see whether the learned result changes with time. If the learning rate is small in the end, we cant decide whether its because the network is stabilized or the learning process is shut down.

**Figure 6:**
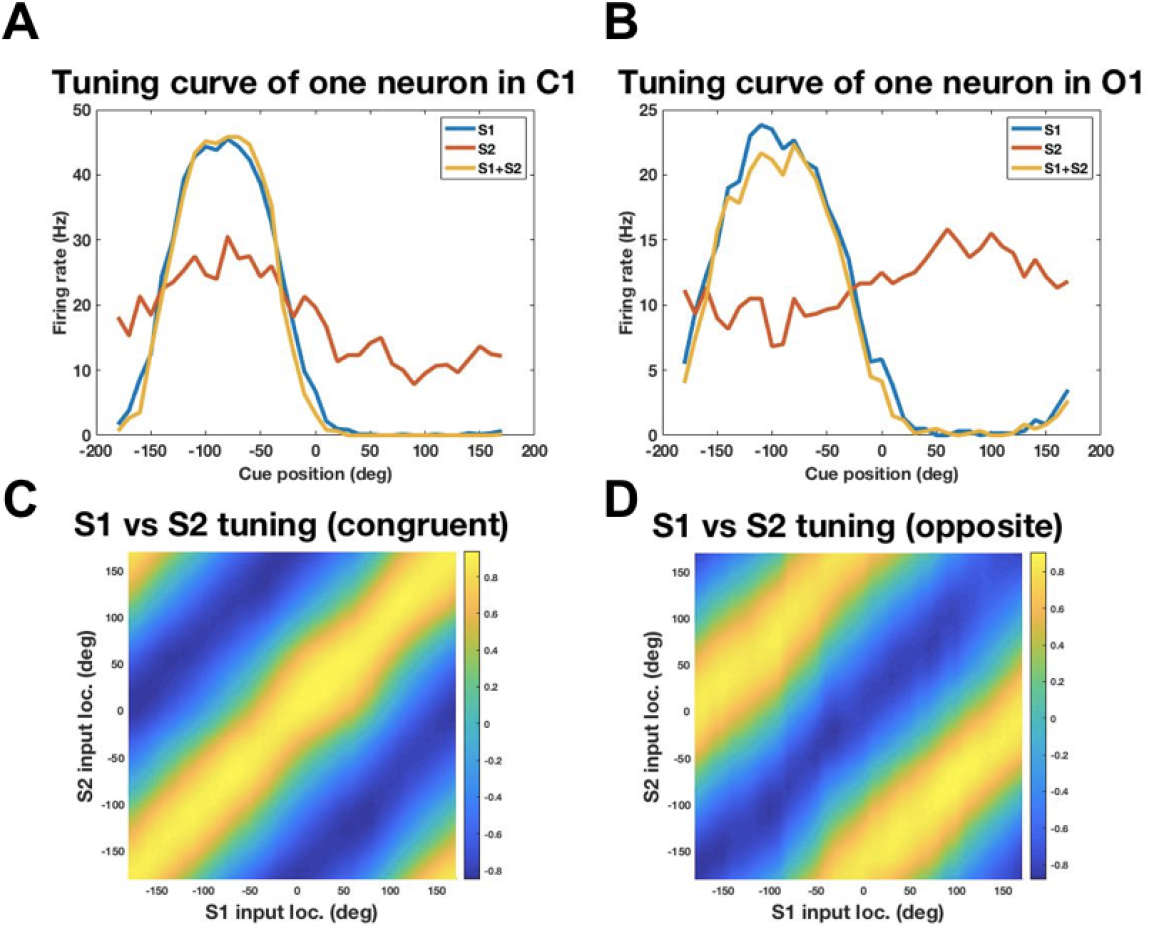
Tuning curves. A) Tuning curve for a congruent neuron preferring −90 degree. B) Tuning curve for an opposite neuron preferring −90 degree in module 1 and 90 degree in module 2. C) Correlation of tuning curves of C1 neurons towards unimodal S1 stimulus and unimodal S2 stimulus. The bright diagonal means that there is strong correlation of response to S1 and S2 inputs from the same location. D) Correlation of tuning curves of O1 neurons towards unimodal S1 stimulus and unimodal S2 stimulus. A bright ridge along the diagonal shifted by 180 degree means there is strong correlation of response to S1 and S2 inputs from opposite locations.

**Figure 7:**
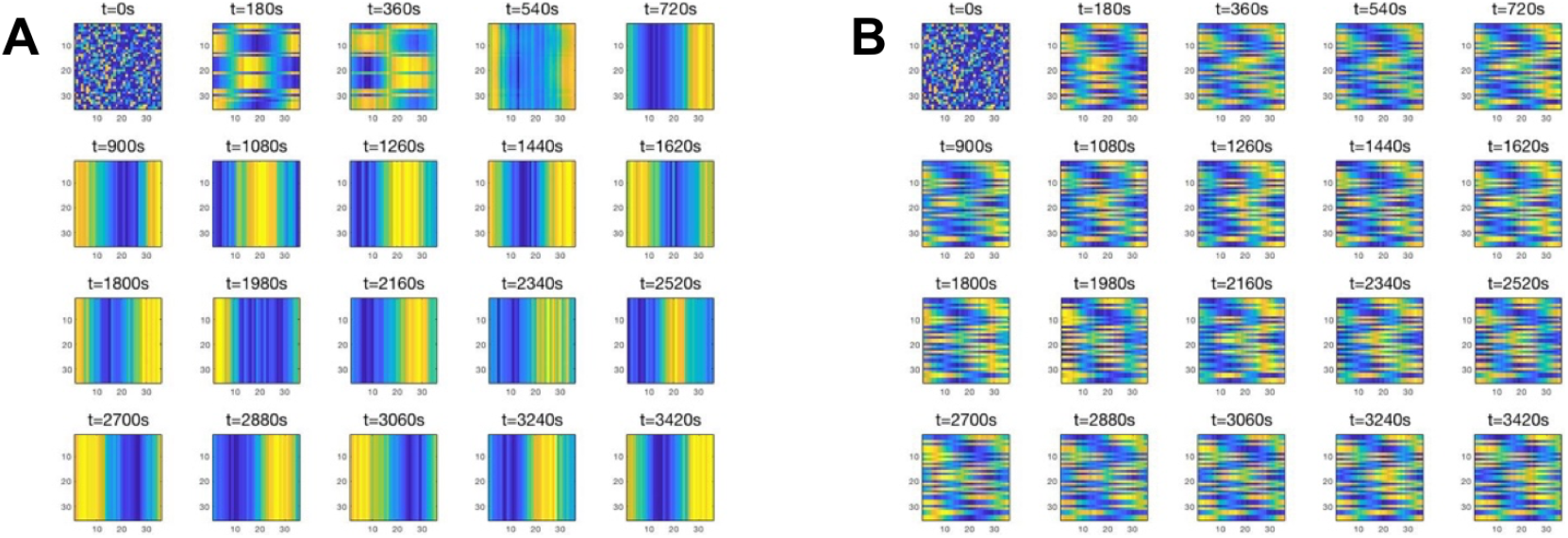
Learned topological diagram’s evolution with time. A) aei=1.2, aie=1.4. This is the case where the inhibition is not strong enough. Every neuron will learn the same thing and it changes with time. B) aei=1.6, aie=1.6. This is the case where the inhibition is too strong. The topological diagram will be messy.

In Figure 8 we demonstrate the parameter dependence of feedforward learning. We vary aei and aie, which are inhibitory to excitatory strength and excitatory to inhibitory strength respectively. The effective inhibition is the convolution of these two parameters.

**Figure 8:**
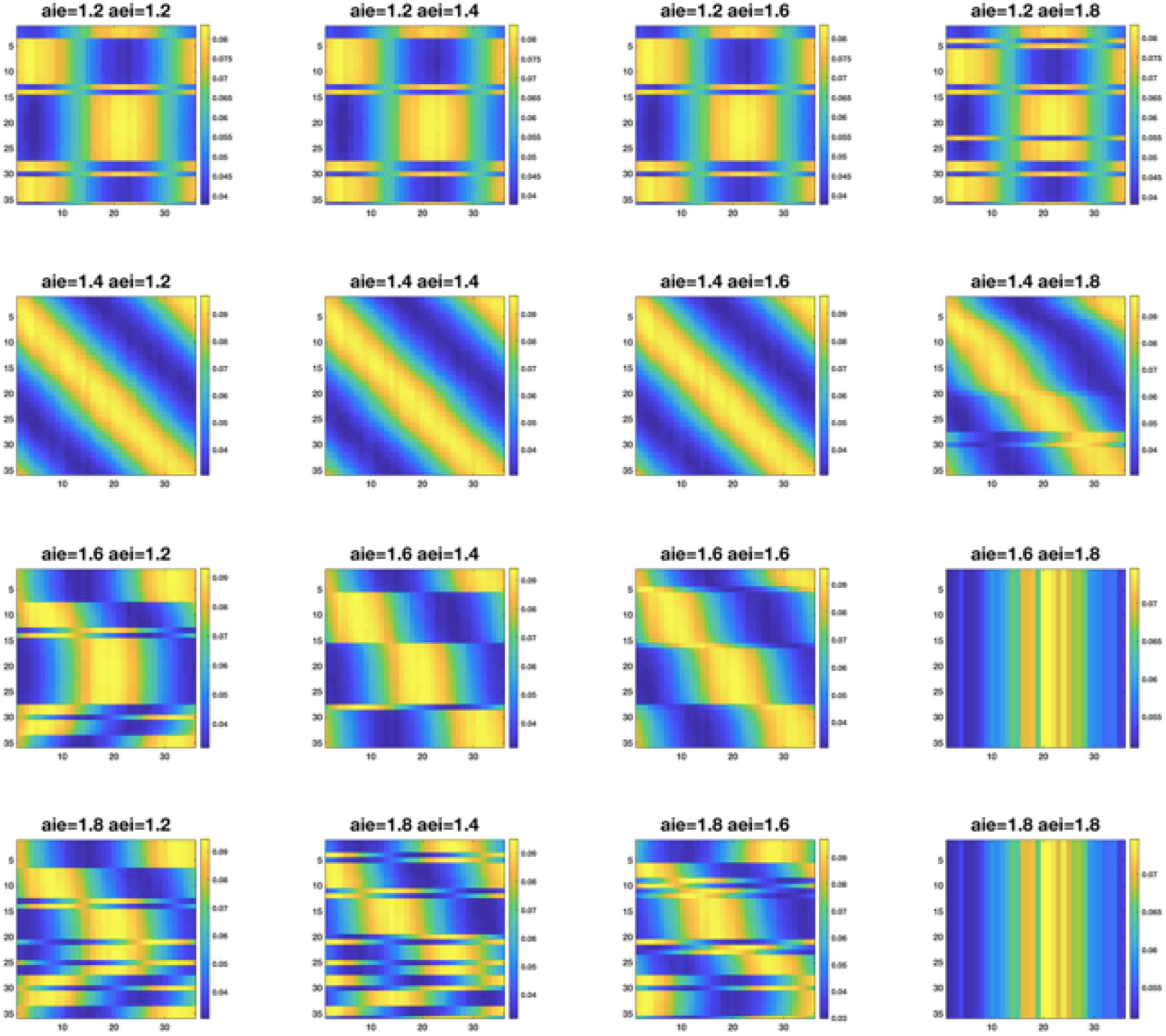
“Phase diagram” of the learned topological diagram. The parameter range is quite narrow.

It is shown in the figure that when the inhibition is too weak, the learned kernels tend to be the same, but not exactly the same, which is because decreased learning rate makes the network trapped in some local minimums. When the inhibition is too strong, the learned topological graph is messy. When the inhibition is much too strong, the learned kernels are all the same again. It has to be made clear that this is due to a numerical inefficiency. When the strengths are that large, the conductance can be so large that the numerical integration step is not small enough. The network behaves abnormally, whose responses get saturated and since the activity of all the neurons are the same, they learn the same kernel.

### 4.2 Inference mode versus learning mode

Parameters in inference mode and learning mode can be quite different. For one thing, the implementation of divisive normalization is critical. We have experimented two kinds of inhibition one is that inhibitory neurons are von-Mises, and the other requires the inhibition to be uniform. Both will work for operation when the connections S1-C1 are learned, but uniform (and with sufficient strength) is needed during learning, otherwise, all the neurons responses will saturate, and receptive fields kept following the input training stimuli. It is also important for each stimulus, there is a clear winner, so that it can suppress the others. A von-Mises inhibition kernel tends to suppress oneself, and if the width is not wide enough, it will allow another winner to emerge at the far away, causing the pool to learn two cycles in the topology, two sets of neurons with the same receptive fields.

For another, during learning there needs to be a weak reciprocal (long range) connection strength so that the learning in two modules wont interfere with each other. However, during inference, the reciprocal connection strength needs to be strong so that congruent neurons and opposite neurons will get enough information from the other module to perform integration.

### 4.3 Decreased learning rate with time

At first, the learned feedforward connections from S to C or O are too noisy to produce a reasonable population response. So we adopted an exponentially decreasing learning rate with time from 0.2 to 0.02 in 100 steps. To comprehend this intuitively, the system may not reach local minimum with too high a learning rate kept unchanged. It may bounce back and force on the energy curve. Actually, at first there needs to be a large learning rate to avoid being caught in local minimum. Later there needs to be a small learning rate to learn the details of the curves.

We have also tried decreasing input noise strength with time, but found it neither sufficient nor necessary. Also, biologically input noise from other cortical areas basically wont change with time.

## 5 Conclusion

Here, we first demonstrate that the decentralized neural circuit model for multisensory processing, a firing rate model without the explicit representation of inhibitory neurons (thus violating Dale’s law), can be implemented by leaky integrate and fire model, originally developed for modeling V1 neurons, wiht realistic parameters. This is simply achieved by (1) introducing inhibitory neurons for performing normalization, i.e., PV neurons (the weight is fixed or parameters to be tuned), and (2) introducing an inhibitory neuron for each C1 to relay the inhibition to the O neurons (Wc1i12 is fixed von-Mise but Wi12o1 is learned). The shape learned is von-Mise. It does have some inhibitory synaptic plasticity. However, we can’t learn both Wc1i12 and Wi12o1 because there might be many options-degenerate. This model is different from Zhang et al.’s in that the opposite neurons can be completely learned in each module without connections from the other module. Essentially, the model can accomplish what firing rate model accomplished, but more biologically realistic in (1) obeying Dale’s law, (2) having spiking activities, following detailed V1 model at cellular molecular level, for example, having AMPA and NMDA receptor dynamics. More interestingly, we find that the model that was developed for V1 can be readily transferred to model MSTd and VIP multi-sensory integration circuit, indicating that this model might be a biologically realistic canonical model. That would imply that the multi-sensory integration circuit might also be relevant for understanding the computation in V1 itself. For example, it is reasonable to conjecture that the congruent neurons and the opposite neurons might also be in V1. That is, the congruent neurons might be integrating information across hypercolumns, while the opposite neurons are measuring the tension and disparity between the information in the different hypercolumns. When the different hypercolumns disagree, opposite neurons will fire strongly, driving the divorce of the two hypercolumns, as in the Mumford-Shah equation in image segmentation.

Second, we showed that such model can be using Willshaw law. We found that the parameters needed for the models, particularly inhibitory spatial range and strength for normalization is more stringent during development (learning) than during operation. For example, during operation, a large Gaussian (von-Mise) inhibitory kernel would be sufficient. But during development, uniform inhibition across all directions are required. There also needs to be delicate balance between inhibition and excitation for this to work.

## Supporting information

source tex file for the paper

---

**Table 1:** Simulation details

| parameter | value | meaning |
| --- | --- | --- |
| $g_L$ | 50 | leaky conductance |
| $V_R$ | 0 | reset potential |
| $V_T$ | 1 | voltage threshold |
| $V_I$ | -2/3 | inhibitory reversal potential |
| $V_E$ | 14/3 | excitatory reversal potential |
| $t_R$ | 3(1)ms | refractory period for excitatory (inhibitory) neurons |
| $\tau_{AMPA}$ | 1ms | time constant for AMPA receptor |
| $\tau_{NMDAr}$ | 2ms | rise time constant for NMDA receptor |
| $\tau_{NMDAd}$ | 128ms | decay time constant for NMDA receptor |
| $\tau_{GABAA}$ | 1.67ms | time constant for GABAA receptor |
| $\tau_{GABAB}$ | 7ms | time constant for GABAB receptor |
| $f_{rtE}$ | 10Hz | excitatory noise's firing rate |
| $f_{rtI}$ | 10Hz | inhibitory noise's firing rate |
| $dt$ | 0.1ms | time step for potential update |
| $t_{final}$ | 3600s | total simulation time |
| $S_{EE}$ | 10.04 | short range excitatory-excitatory synaptic strength |
| $S_{EI}$ | 1037.89 | short range inhibitory-excitatory synaptic strength |
| $S_{IE}$ | 181.38 | short range excitatory-inhibitory synaptic strength |
| $L_{EE}$ | 1.408 | long range excitatory-excitatory synaptic strength |
| $a_{IE}$ | 1.5 | uniform kernel for excitatory pool to inhibitory normalization pool |
| $a_{EI}$ | 1.5 | uniform kernel for inhibitory normalization pool to excitatory pool |
| $S_E$ | 15000 | synaptic strength of excitatory noise |
| $S_I$ | 10000 | synaptic strength of inhibitory noise |
| $J^{rc1}$ | 8 | controlling amplitude of recurrent connection (module 1) |
| $J^{rc2}$ | 8 | controlling amplitude of recurrent connection (module 2) |
| $J^{rp}$ | 8 | controlling amplitude of recurrent connection between C1 and C2 |
| $\kappa_1$ | 3 | controlling width of recurrent connection connection in module 1 |
| $\kappa_2$ | 3 | controlling width of recurrent connection connection in module 2 |
| $a_1$ | 0.8 | controlling width of input in module 1 |
| $a_2$ | 0.8 | controlling width of input in module 2 |
| $k_1$ | 200 | controlling amplitude of input in module 1 |
| $k_2$ | 200 | controlling amplitude of input in module 2 |
| $\alpha$ | 0.5 | controlling decay term in Willshaw's rule |
| $dt2$ | 50ms | time step for connection weight update |
| $\Delta t$ | 100ms | time window for learning |

