## Supplementary material for "A spiking neural circuit model for learning multi-sensory integration": source tex file for the paper: ref.bib

@book{stein2004handbook,
  title={The Handbook of Multisensory Processes},
  author={Stein, G.C.C.S.B.E. and Calvert, G. and Spence, C. and Spence, D.E.P.C. and Stein, B.E. and Stein, P.C.B.E.},
  isbn={9780262033213},
  lccn={2004042612},
  series={A Bradford book},
  url={https://books.google.com/books?id=CZS\_yDoFV7AC},
  year={2004},
  publisher={MIT Press}
}

@book{spence2004crossmodal,
  title={Crossmodal Space and Crossmodal Attention},
  author={Spence, C. and Driver, J. and Driver, J.C.},
  isbn={9780198524861},
  lccn={2004302311},
  url={https://books.google.com/books?id=6UceYiqZGg0C},
  year={2004},
  publisher={OUP Oxford}
}

@article{neural_correlates,
author = {Gu, Yong and Angelaki, Dora and Deangelis, Gregory},
year = {2008},
month = {10},
pages = {1201-10},
title = {Neural correlates of multisensory cue integration in macaque MSTd},
volume = {11},
journal = {Nature neuroscience},
doi = {10.1038/nn.2191}
}

@article{representation,
author = {Chen, Aihua and Deangelis, Gregory and Angelaki, Dora},
year = {2011},
month = {08},
pages = {12036-52},
title = {Representation of Vestibular and Visual Cues to Self-Motion in Ventral Intraparietal Cortex},
volume = {31},
journal = {The Journal of neuroscience : the official journal of the Society for Neuroscience},
doi = {10.1523/JNEUROSCI.0395-11.2011}
}

@article{functional,
author = {Chen, Aihua and Deangelis, Gregory and Angelaki, Dora},
year = {2013},
month = {02},
pages = {3567-81},
title = {Functional Specializations of the Ventral Intraparietal Area for Multisensory Heading Discrimination},
volume = {33},
journal = {The Journal of neuroscience : the official journal of the Society for Neuroscience},
doi = {10.1523/JNEUROSCI.4522-12.2013}
}

@article{visual_and,
author = {Gu, Yong and Watkins, Paul and Angelaki, Dora and Deangelis, Gregory},
year = {2006},
month = {02},
pages = {73-85},
title = {Visual and nonvisual contributions to three-dimensional heading selectivity in the medial superior temporal area},
volume = {26},
journal = {The Journal of neuroscience : the official journal of the Society for Neuroscience},
doi = {10.1523/JNEUROSCI.2356-05.2006}
}

@article{multisensory,
author = {Morgan, Michael and Deangelis, Gregory and Angelaki, Dora},
year = {2008},
month = {09},
pages = {662-73},
title = {Multisensory Integration in Macaque Visual Cortex Depends on Cue Reliability},
volume = {59},
journal = {Neuron},
doi = {10.1016/j.neuron.2008.06.024}
}

@article{complementary,
author = {Zhang, Wen and Wang, He and Chen, Aihua and Gu, Yong and Lee, Tai and Wong, K. Y. Michael and Wu, Si},
year = {2019},
month = {05},
pages = {},
title = {Complementary congruent and opposite neurons achieve concurrent multisensory integration and segregation},
volume = {8},
journal = {eLife},
doi = {10.7554/eLife.43753}
}

@article{emergence,
author = {Niu, Xueyan and Chau, Ho and Lee, Tai and Zhang, Wen},
year = {2019},
month = {11},
pages = {},
title = {Emergence of opposite neurons in a decentralized firing-rate model of multisensory integration},
doi = {10.1101/845743}
}

@article{orientation,
author = {Tao, Louis and Cai, David and McLaughlin, David and Shelley, Michael and Shapley, Robert},
year = {2006},
month = {09},
pages = {12911-6},
title = {Orientation selectivity in visual cortex by fluctuation-controlled criticality},
volume = {103},
journal = {Proceedings of the National Academy of Sciences of the United States of America},
doi = {10.1073/pnas.0605415103}
}

@article{modeling,
author = {Rangan, Aaditya and Cai, David and McLaughlin, David},
year = {2006},
month = {01},
pages = {18793-800},
title = {Modeling the spatiotemporal cortical activity associated with the line-motion illusion in primary visual cortex},
volume = {102},
journal = {Proceedings of the National Academy of Sciences of the United States of America},
doi = {10.1073/pnas.0509481102}
}
