## Supplementary figures and images for "A spiking neural circuit model for learning multi-sensory integration"

### 1.png

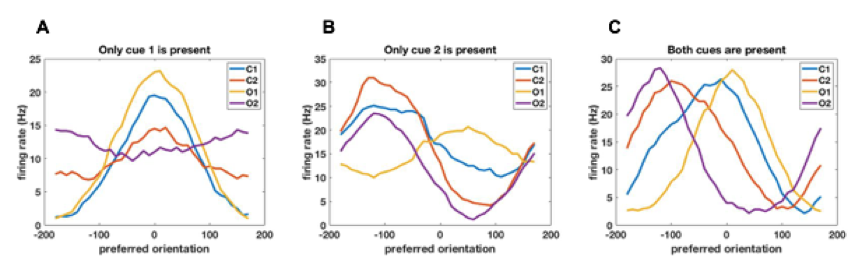

### 2.jpg

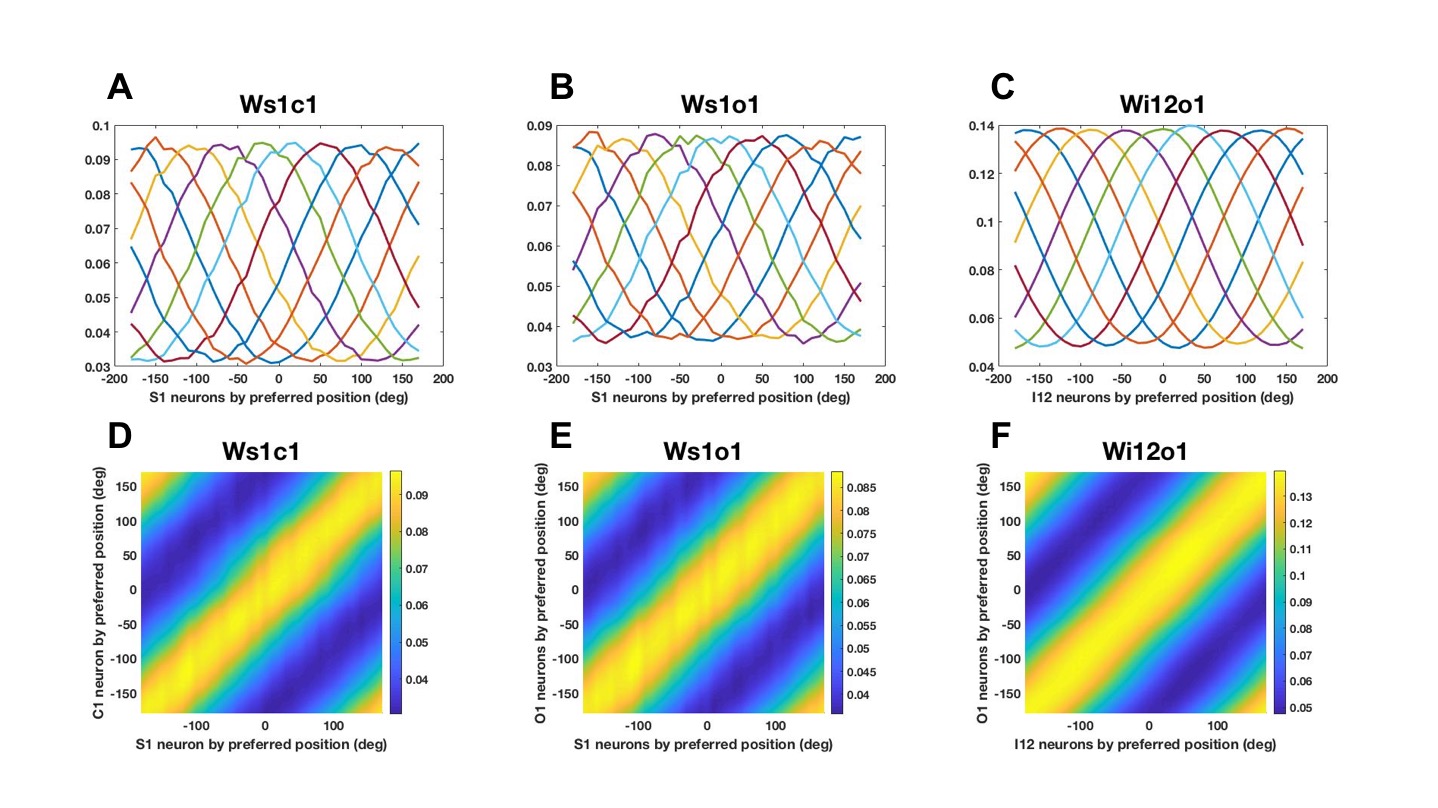

### 3.jpg

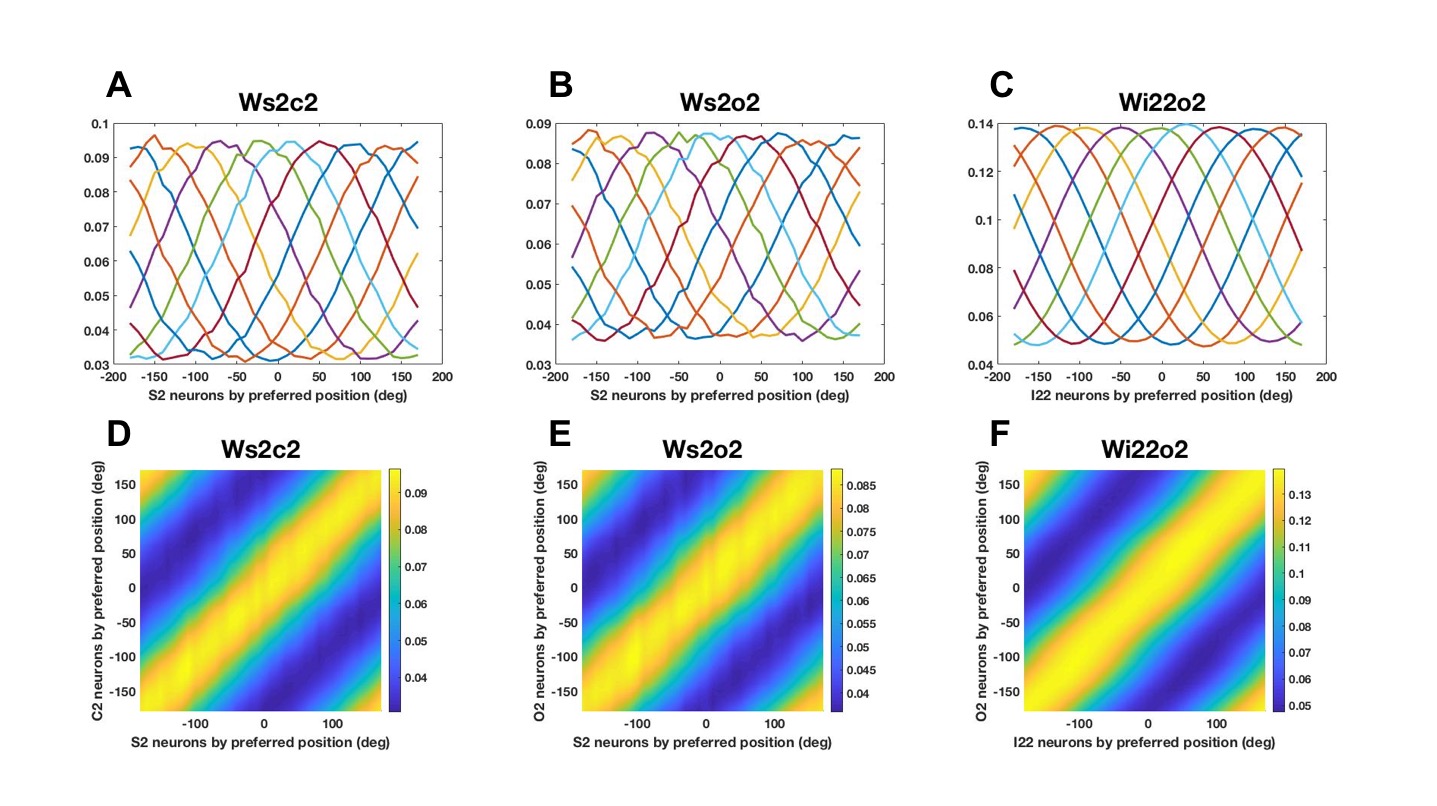

### 4.jpg

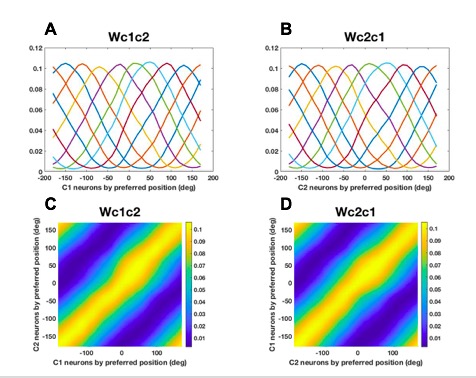

### 5.png

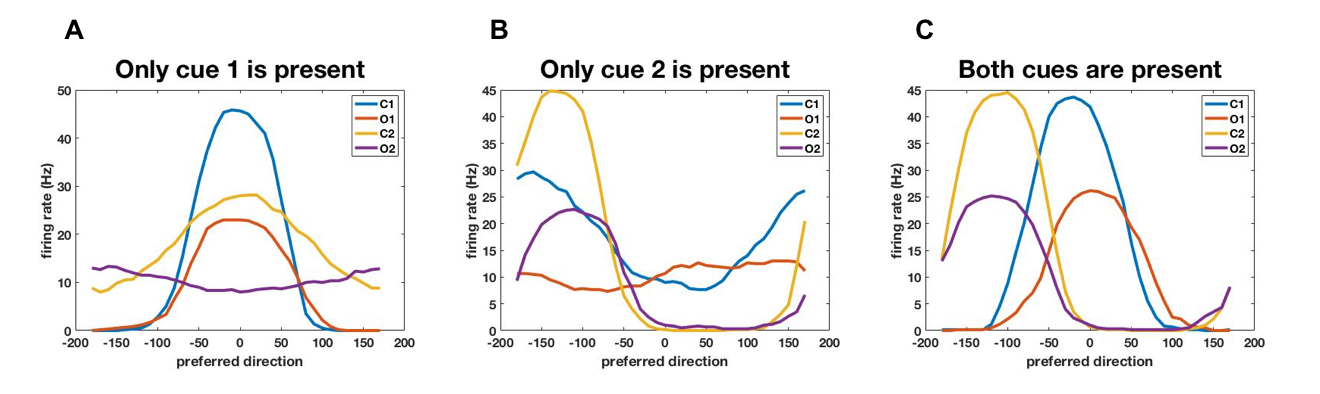

### 6.png

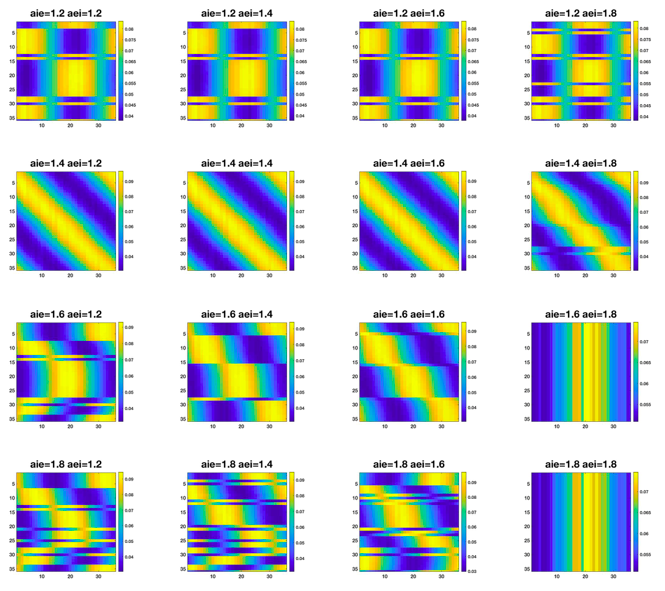

### 7.png

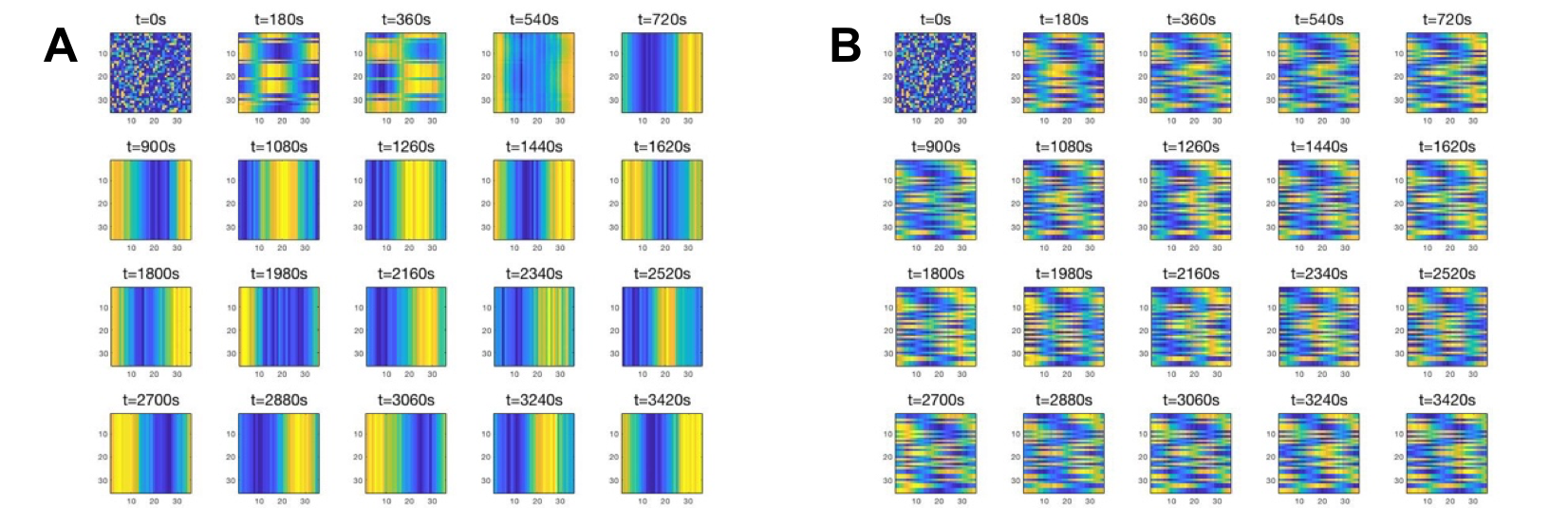

### o05271.png

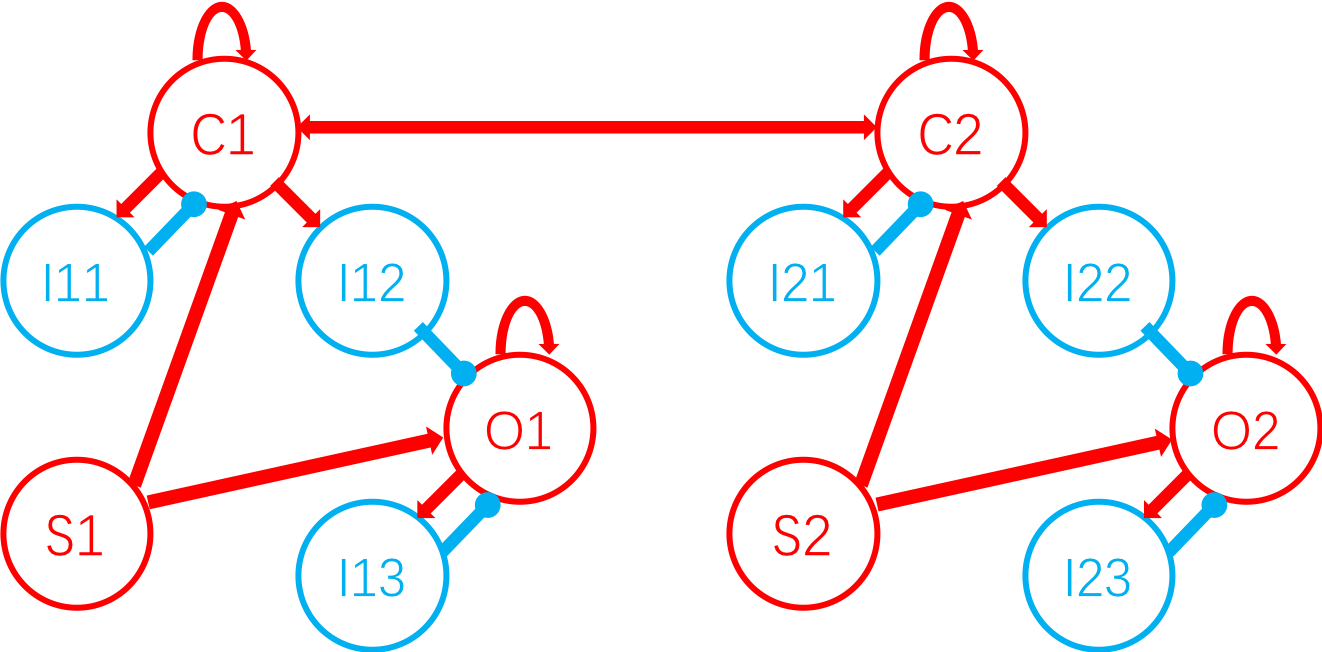

### o10061.png

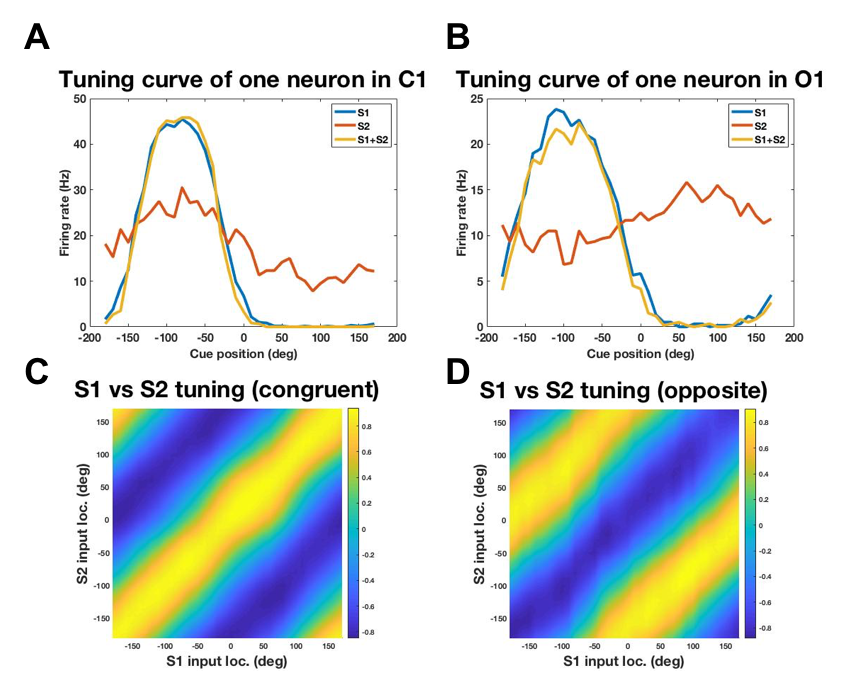

### o11176.jpg

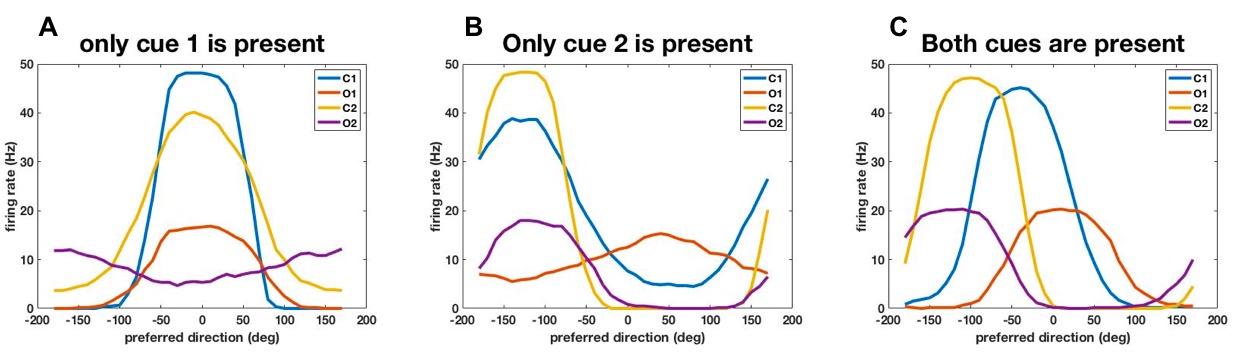
